# A Field-Deployable Diagnostic Assay for the Visual Detection of Misfolded Prions

**DOI:** 10.1101/2021.11.22.469560

**Authors:** Peter R. Christenson, Manci Li, Gage Rowden, Marc Schwabenlander, Tiffany M. Wolf, Sang-Hyun Oh, Peter A. Larsen

## Abstract

Chronic Wasting Disease (CWD), a prion disease of cervids, has been identified across North America, Northern Europe and Korea. Diagnostic tools for the rapid and reliable detection of prion diseases are limited. Here, we combine gold nanoparticles (AuNPs) and quaking induced conversion (QuIC) technologies for the visual detection of amplified misfolded prion proteins from tissues of wild white-tailed deer infected with Chronic Wasting Disease (CWD). Our newly developed diagnostic test, MN-QuIC, enables both naked-eye and light-absorbance measurements for the detection of misfolded prions. The MN-QuIC assay leverages basic laboratory equipment that is cost-effective and portable, thus facilitating real-time prion diagnostics across a variety of settings. To test the portability of our assay, we deployed to a rural field station in southeastern Minnesota and tested for CWD on site. We successfully demonstrated that MN-QuIC is functional in a non-traditional laboratory setting by performing a blinded analysis in the field and correctly identifying all CWD positive and CWD not detected (independently confirmed with ELISA and/or IHC tests) animals at the field site, thus documenting the portability of the assay. Additionally, we show that electrostatic forces and concentration effects help govern the AuNP/prion interactions and contribute to the differentiation of CWD-prion positive and negative samples. We examined 17 CWD-positive and 24 CWD-not-detected white-tailed deer tissues that were independently tested using ELISA, IHC, and RT-QuIC technologies, and results secured with MN-QuIC were 100% consistent with these tests. We conclude that hybrid AuNP and QuIC assays, such as MN-QuIC, have great potential for sensitive, field-deployable diagnostics for a variety of protein misfolding diseases.

## Introduction

A common feature of many neurodegenerative diseases is the presence of misfolded proteins that accumulate within the central nervous system, ultimately contributing to advanced neurodegeneration and death. Misfolded protein diseases (proteopathies) impact a wide variety of mammals, including Creutzfeldt-Jakob disease (CJD), Alzheimer’s disease, and Parkinson’s disease in humans, bovine spongiform encephalopathy (BSE) in cattle, scrapie in sheep, pituitary pars intermedia dysfunction in horses and chronic wasting disease (CWD) in cervids.^1–5^ Given well-documented diagnostic limitations surrounding proteopathies in both animals and humans, (i.e., poor sensitivity, limited antibodies for immuno-based assays, etc.), it is imperative to develop improved diagnostic assays.^2,6–11^ With respect to prion diseases (a proteopathy subclass caused by infectious proteins), tests that could be deployed in a variety of settings (e.g., hospitals, veterinary clinics, field stations, etc.) would greatly aid the detection of infectious prions thus limiting their spread. It is within this framework that we approach the development of diagnostic tools for CWD of cervids, a model neurodegenerative disorder with urgent needs for portable diagnostic assays that would facilitate rapid detection, thus preventing CWD prions from entering human and animal food-chains.

Similar to CJD in humans, which rapidly progresses and is always fatal, and BSE in cattle (commonly known as “mad cow disease”), CWD is a prion disease or Transmissible Spongiform Encephalopathy (TSE) that is 100% fatal to infected animals and has no treatments or vaccines.^1,12^ CWD impacts cervids across North America, Scandinavia, and South Korea^13,14^ (e.g., elk, moose, mule deer, white-tailed deer, reindeer). The disease continues to spread to new cervid populations, and there are increasing health concerns for both humans and animals exposed to various CWD prion strains.^13–15^ All mammals have native functioning cellular prion protein distributed throughout various tissues and playing essential roles in a variety of physiological functions, especially those of the central nervous system.^16^ Like other protein misfolding neurodegenerative disorders, native prions in cervids adopt pathogenic conformations of misfolded prions (PrP^CWD^). PrP^CWD^ propagates throughout an infected animal, forming fibrils that accumulate in lymph and nervous tissues, leading to death years after exposure (Fig. 1a). CWD poses obvious risks to the health of impacted cervid populations globally, and the disease is an immediate threat to not only cervid health, but also all cervid-related economies. Indeed, cervid hunting and related activities generate tens of billions of USD annually in the United States.^17^ Across the world, cervids provide a wide array of economically/medicinally^18^ important products that are routinely consumed and/or used by humans (i.e., venison meat, antler velvet health supplements, antlers, hides, etc.).

**Figure 1.**
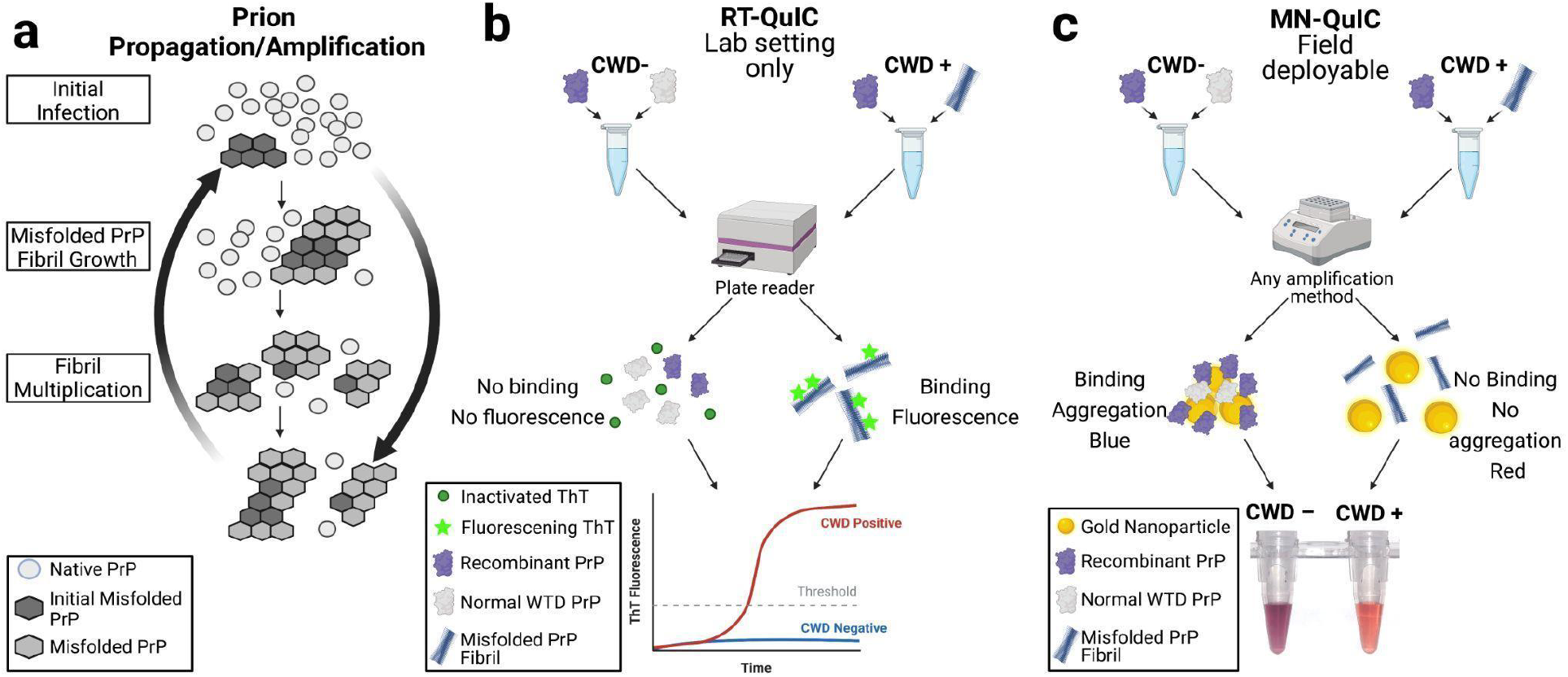
Overview of the amplification assays. **a.)** schematic of misfolded fibril growth. The initial PrP^CWD^ causes native PrP to misfold. **b.)** Overview of RT-QuIC. PrP^CWD^ seeds originating from tissue samples of cervids are added to rPrP solutions. These solutions are then shaken and incubated for ∼48 hrs. If present, PrP^CWD^ induces conformational changes of the rPrP. Amplification results are read in real-time with ThT fluorescence. **c.)** MN-QuIC overview. After 24 hr amplification, products are diluted and added to an AuNP solution. CWD positive samples result in a red solution (peak absorbance wavelength ∼516 nm) while CWD negative solutions are purple (peak absorbance wavelength ∼560 nm).

Current diagnostic methods for the detection of protein-misfolding diseases, including CWD and other TSEs, are limited.^2,6–11^ Commonly utilized TSE diagnostic assays rely heavily on antibody-based enzyme-linked immunosorbent assay (ELISA) and immunohistochemistry (IHC) technologies that are expensive, time-consuming, and require substantial training and expertise to operate.^6^ A feature of ELISA and IHC assays is that the antibodies routinely used cannot differentiate between native PrP and misfolded TSE-associated prion proteins (PrP^TSE^) thus requiring enzymatic digestion to enrich for PrP^TSE^, a methodology that may impact diagnostic sensitivity through the destruction of particular TSE-affiliated PrP strains.^19,20^ Collectively, these antibody-based assays are limited in the identification of early-stage TSE infections, and they are primarily used on tissues collected post-mortem.

The detection of prion seeding activity was recently enhanced by various assays involving the amplification of protein misfolding *in vitro*, including protein misfolding cyclic amplification (PMCA), ^6,21^ end-point quaking-induced conversion (EP-QuIC)^22,23^ and real-time quaking-induced conversion (RT-QuIC).^6,24–27^ Of these prion amplification methods both EP-QuIC and RT-QuIC (Fig. 1b) utilize a recombinant mammalian PrP substrate (rPrP) that is incubated and shaken with the diagnostic samples. When PrP^TSE^ is present within a given QuIC reaction, it induces a conformational change of the rPrP, forming a beta-sheet enriched mixture that is quantified with fluorescent Thioflavin T (ThT) measurements. Despite the advantages of EP-QuIC and RT-QuIC, there still remain major limitations, including the need for expensive and large laboratory equipment and complex strategies for visualizing and analyzing results. This limits these methods to well-funded research laboratories or large diagnostic labs. In short, more effective TSE diagnostic methods that leverage small and portable equipment with straightforward results are needed to rapidly detect various TSEs in widespread surveillance and prevent additional spread and introduction of TSE prions into human and animal food chains. This is especially true for CWD, as the disease continues to expand across both farmed and wild cervid populations.

In parallel to diagnostic advancements based on prion and protein amplification methods, gold nanoparticles (AuNPs) have been increasingly used for medical applications including disease diagnostics,^28–30^ drinking water safety, ^31^ and food safety. ^32^ Due to localized surface plasmon resonances (density fluctuation of conduction electrons)^33^, AuNPs have optical absorption peaks that are sensitive to the environment at the AuNP’s surface. ^33–37^ These unique optical properties make plasmonic nanoparticles useful in color-based detection assays. ^38,39^ Previous studies have indicated affinities of prion proteins with a variety of bare and functionalized metals, including gold.^39–42^ One limitation of AuNPs is that they are susceptible to nonspecific binding.^43^ Real biological samples are not homogeneous and have many different proteins, ions, and other organic molecules associated with them that can induce AuNPs to spontaneously aggregate making effective AuNP diagnostics challenging. To overcome this, we sought to demonstrate the diagnostic utility of gold nanoparticles for detecting misfolded PrP^CWD^ within QuIC amplified products of CWD positive and negative white-tailed deer (WTD) tissues.

By combining the unique plasmonic properties of AuNPs and the methods of quaking-based prion protein fibril amplification, we have successfully overcome the non-specific binding challenges associated with AuNP diagnostics in real samples and have produced a nanoparticle-based assay (herein named Minnesota-QuIC; MN-QuIC) that can detect the presence or absence of misfolded PrP^CWD^ using both visual and spectroscopic methods (Fig. 1c). This method uses only basic lab equipment which allows it to be deployed outside the laboratory. In March of 2021, we deployed the MN-QuIC assay to a field station in rural southeastern Minnesota where the Minnesota Department of Natural Resources (DNR) was performing its annual CWD surveillance and targeted culling efforts. We demonstrated proof of concept experiments for MN-QuIC’s utility as a portable prion assay by successfully detecting CWD-infected WTD tissues at the DNR field station.

## Results

### Gold nanoparticle interaction with native cellular prions vs. misfolded fibrils

To investigate whether AuNPs can differentiate between misfolded PrP fibrils and native PrP originating from recombinant hamster prion protein (rPrP), two sets of reaction mixtures seeded with and without spontaneously misfolded rPrP prion fibrils were processed following modified RT-QuIC protocols without ThT. ^44,45^ The presence of fibril formation was examined in all reaction mixtures by adding and quantifying ThT *post hoc* (Fig. S1a). ThT fluorescence was significantly different between misfolded rPrP (seeded) and native non-misfolded rPrP (no seed) (p=0.05; Fig. S1a). We hypothesized that misfolded prion fibrils would interact differently with AuNPs, as compared with native rPrP, and that the interaction would influence AuNP aggregation as measured by dynamic light scattering (DLS). Following ThT quantification, misfolded rPrP samples and native rPrP samples were spiked into separate AuNP solutions. After a 30 min incubation period at ambient temperature, there was a visible color difference between the AuNP solutions spiked with misfolded and native rPrP similar to the change in figure 1c. This was also reflected in the visible spectrum of the absorbance peak (Fig. 2a).

**Figure 2.**
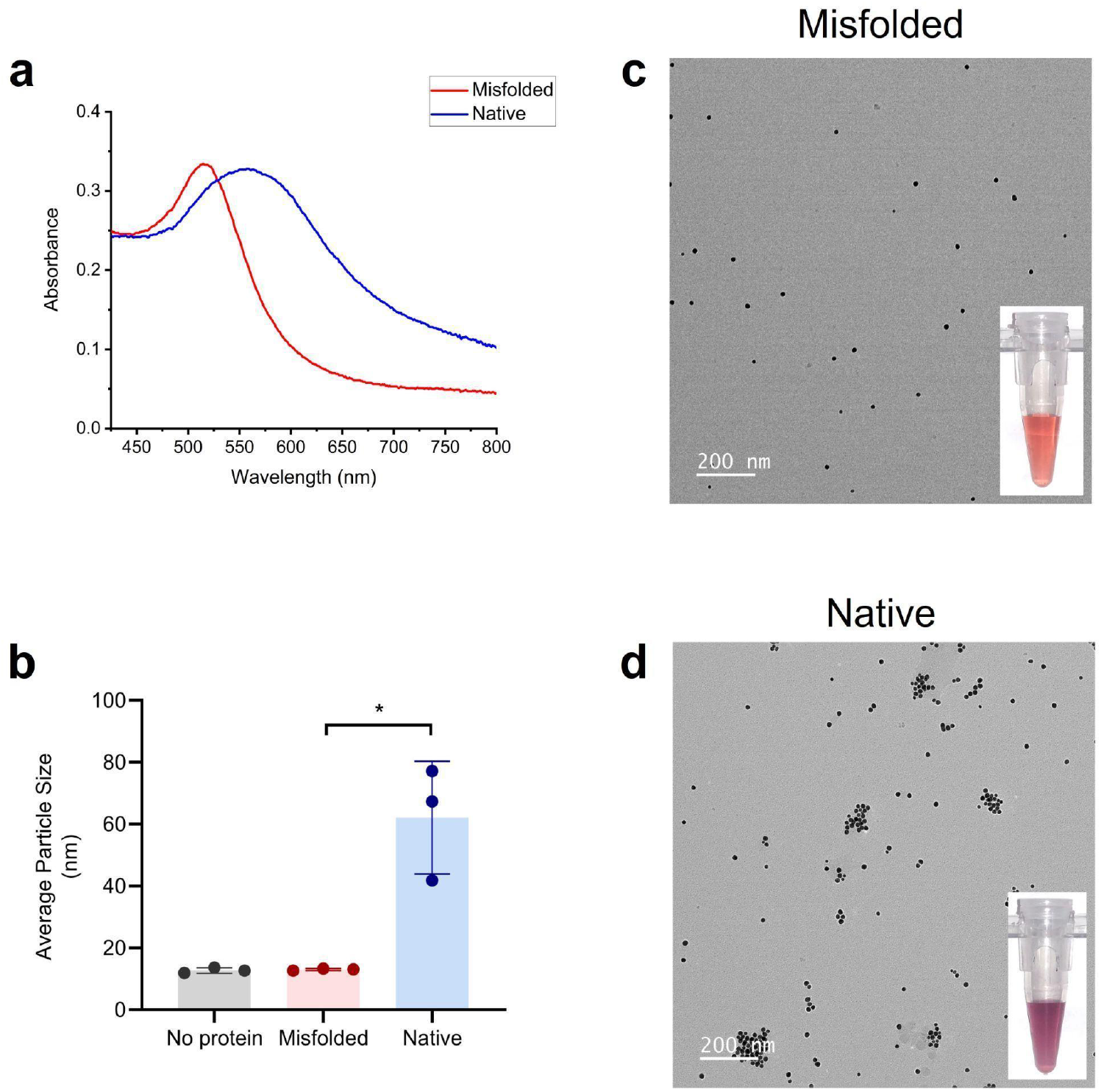
**a**. The absorbance spectrum of AuNP solutions spiked with misfolded rPrP from seeded QuIC reactions (red line) and non-misfolded/native rPrP from QuIC reactions without seed (blue line). **b**. Average effective particle sizes in AuNP solutions containing no protein, misfolded rPrP, and native rPrP observed by DLS. **c**. TEM image of 15nm AuNPs after being spiked with post amplified misfolded PrP. Inset shows an example of AuNP solution after being spiked with misfolded PrP. **d**. TEM image aggregating AuNPs after being spiked with native PrP. Inset shows an example of AuNP solution after being spiked with post amplified native PrP. *, p-value < 0.05, error bars show standard deviation.

Color changes due to aggregation have been reported in the literature for a variety of nanoparticle/protein combinations including prions and AuNPs. ^38,39^ To quantify this, DLS experiments were performed on three AuNP solutions (seeded, non-seeded, and blank; see Methods), and average effective particle size was determined for each sample (Fig. 2b). We observed a significant difference of AuNP effective particle sizes between AuNP solutions seeded with misfolded rPrP versus native non-seeded rPrP (p=0.05) (Fig. 2b). The AuNPs spiked with no protein (blank) and misfolded rPrP exhibited similar particle size distributions (Fig. S1b,c), indicating that the misfolded rPrP solutions did not induce aggregation. On the contrary, the addition of diluted native rPrP resulted in larger particle sizes for AuNPs (Fig. S1d) than AuNPs with no protein added (Fig. S1b), indicating that the addition of diluted native rPrP caused AuNPs to aggregate. To further examine this, AuNP solutions spiked with spontaneously misfolded rPrP and native rPrP were studied in a transmission electron microscope (TEM). Through TEM studies it is clear that the AuNPs did not aggregate when spiked with misfolded prion (Fig 2c). Conversely, it can be seen that AuNPs aggregate in the presence of native rPrP (Fig 2d). These results match well with DLS measurements and indicate differential AuNP binding interaction between native rPrP and misfolded rPrP fibrils.

### CWD positive and negative samples produce unique AuNP optical signatures

Understanding that pathogenic prions can induce rPrP misfolding and amplification ^46^ and thus influence AuNP aggregation (see above), we then investigated the potential of MN-QuIC for CWD diagnostics using misfolded PrP^CWD^ positive and negative WTD lymphoid tissues. We used homogenates of independently confirmed CWD positive and negative WTD medial retropharyngeal lymph nodes (RPLN) (Table S1). Independent RT-QuIC analyses were performed on all tissues used for AuNP analyses (Fig. 3a). ^27^

**Figure 3.**
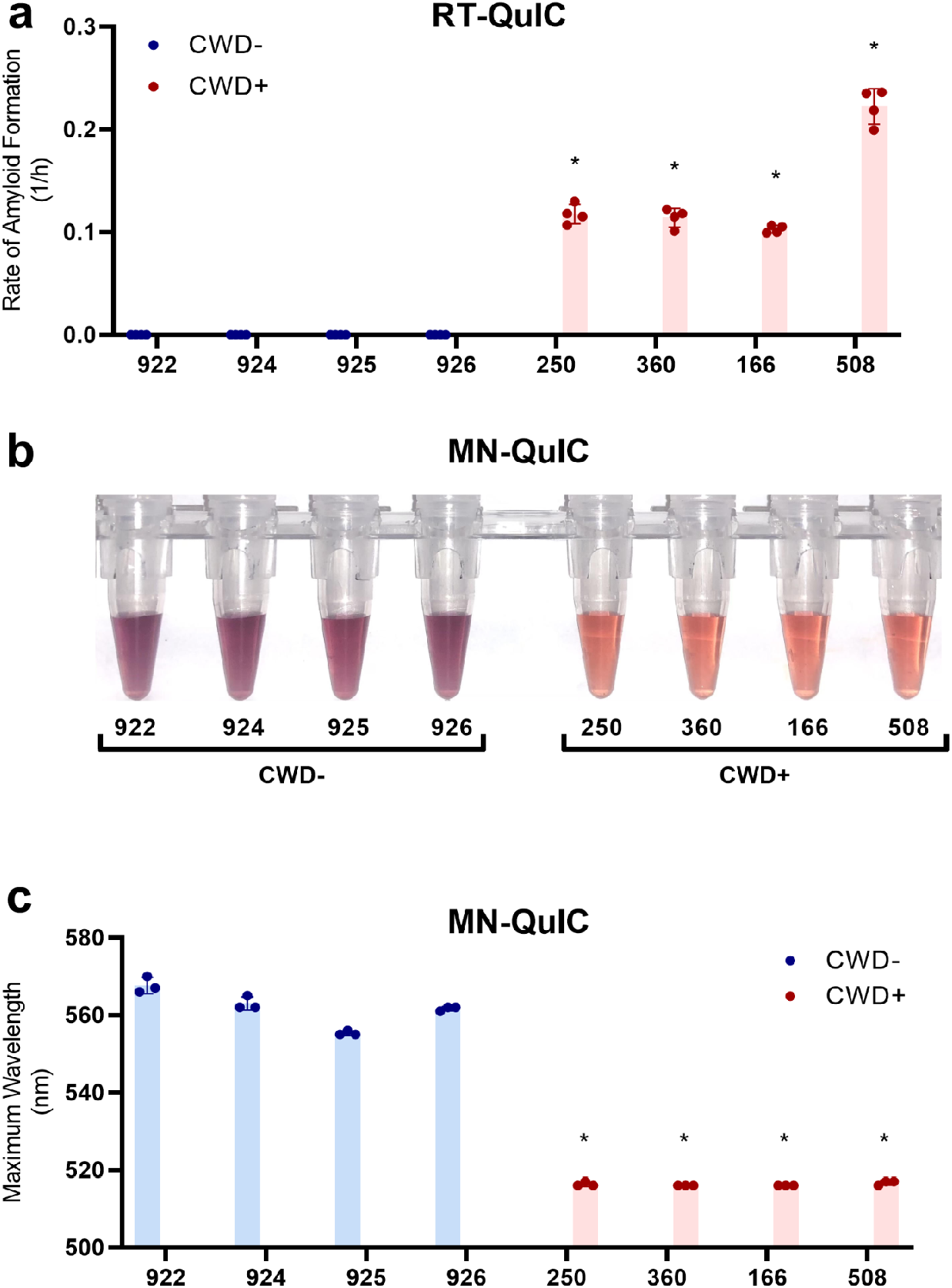
Comparison of RT-QuIC and MN-QuIC results for CWD positive and negative tissues. Sample identification number on the horizontal axis. **a**. RT-QuIC data for the rate of amyloid formation for negative and positive medial retropharyngeal lymph node tissue samples from wild white-tailed deer using ThT fluorescence. **b**. Photo of MN-QuIC tubes showing the color difference for the same set of tissue samples used in panel a. **c**. MN-QuIC peak absorbance wavelength of the same set of solutions used in panel b. *, p-value < 0.05, error bars show standard deviation.

In light of our previous results, we anticipated that AuNPs could be utilized to facilitate direct visualization of QuIC-amplified misfolded rPrP solutions that were seeded with CWD positive tissue using a standard bench-top thermomixer. Thermomixers have been used previously in conjunction with end-point ThT readings (i.e., EP-QuIC) to determine the presence of CJD prion seeding activity. ^23^ To test this hypothesis, the same set of samples was added to the RT-QuIC master mix without ThT on 96-well plates, which were then subjected to shaking/incubation cycles on a thermomixer for 24 hrs. The post-amplification solutions were then diluted to 50% and added to AuNP solutions. We found that we were able to clearly distinguish CWD positive and negative samples simply through color difference appreciable by naked eye; the QuIC-amplified CWD positive and negative samples were red and purple, respectively (Fig. 3b).

To quantify our observations and measure statistical differences, the absorbance spectrum of the AuNPs was measured from 400-800 nm using a 96-well plate reader. In the resulting absorbance spectrum, AuNP solutions combined with QuIC-amplified CWD positive samples had absorbance peaks near 516 nm (Fig. 3c), similar to the 515 nm absorbance peak of the AuNPs prior to the addition of protein solutions. However, the negative sample absorbance peaks were shifted to longer wavelengths of approximately 560 nm (Fig. 3c), confirming that the purple color of AuNP solutions from QuIC products originating from CWD negative tissue samples was consistent with the observed purple color of AuNP aggregates associated with native rPrP (Fig 2b & Fig S1d). Accordingly, the peak AuNP absorbance wavelengths of CWD negative samples are significantly larger (p<0.05) than CWD-positive samples (Fig. 3c).

### Electrostatic forces and rPrP concentration play a role in AuNP CWD detection

Considering the results described above, we aimed to determine the mechanism underlying AuNP aggregation caused by native rPrP solutions. Studies with prions and other proteins have shown that electrostatic forces help govern the interactions between nanoparticles and proteins. ^39,47,48^ Because the theoretical isoelectric point (IP) of our rPrP is around pH 8.9, ^49^ rPrP is positively charged in the pH 7.4 AuNP buffer whereas citrate capped AuNPs are negatively charged even at pHs well below our buffer. ^50^ Thus at pH 7.4, there exists an electrostatic attractive force between AuNPs and native rPrP that contributes to their interactions (aggregation and the color change). The charge on the protein changes when the pH of the environment is altered and thus the interaction between AuNP and rPrP is disrupted. When the pH of a solution is raised closer to the IP of the rPrP, the charge of the protein will become closer to neutral, decreasing the force of attraction between AuNP and rPrP. We showed that as the pH of the AuNP solution was raised closer to the IP of native rPrP, the absorbance peak of the AuNP-rPrP solution decreased from 530 nm (Fig. 4) while the control AuNP solution with no protein had very little peak deviation from 515 nm. This indicates that electrostatic interactions were partially responsible for facilitating the native rPrP interactions with AuNPs. QuIC-amplified PrP^CWD^ products, on the other hand, have experienced major conformational changes from their native form (as confirmed by ThT beta-sheet binding; Fig. 3a) and have formed fibrils. These fibrils can further clump together to form large aggregates reducing the effective concentration of rPrP in the reaction. When there is a low concentration of free-floating rPrP in the solution, the interaction between prions and AuNPs drops below detectable levels (Fig. S2). Because of the concentration effects from fibril formation and aggregation, the interaction between positive (misfolded) prions and AuNPs is unaltered by pH (Fig. 4).

**Figure 4.**
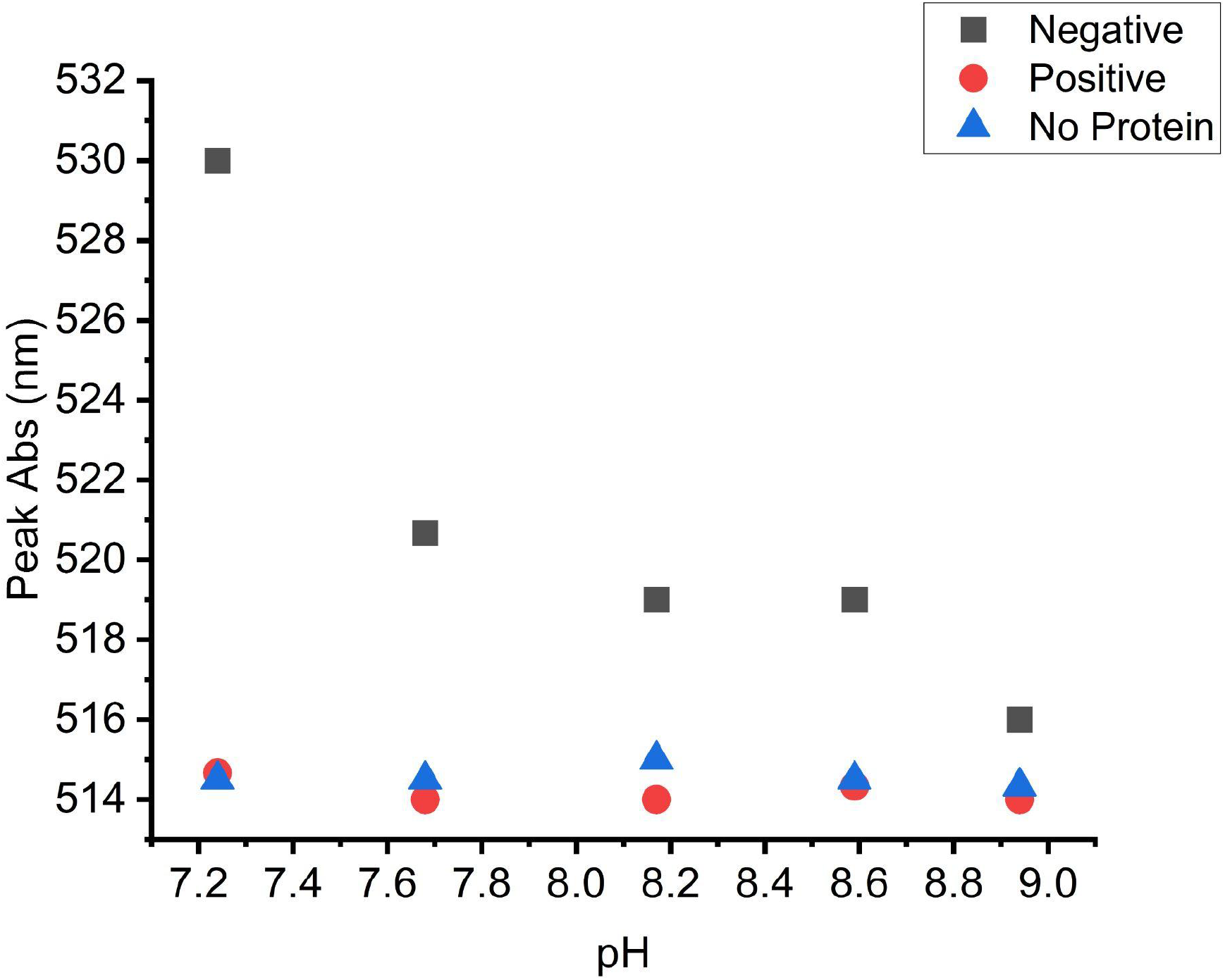
Wavelength of peak absorbance of AuNPs in varying pH buffers containing native CWD negative rPrP (square), PrP^CWD^ positive (circle), and blank/no protein (triangle) solutions.

### Field deployment and higher throughput protocols

To show the potential for a portable, field-deployable diagnostic we performed proof of concept experiments at a rural Minnesota DNR field station (Fig 5a). We tested both pooled and individual tissues consisting of medial retropharyngeal lymph nodes, parotid lymph nodes, and palatine tonsils tissues from 13 WTD that the DNR had recently collected from the surrounding wild deer population. Three of these animals (blinded to the field team) were CWD positive as determined by regulatory ELISA and IHC testing of medial retropharyngeal lymph nodes. Using a blinded testing approach, MN-QuIC successfully detected, via visibly red AuNP solutions (Fig 5b), all three CWD positive animals. We also secured CWD not detected or negative results (purple AuNP solutions) for the 10 animals that were independently identified by ELISA as CWD not detected (as determined by regulatory ELISA testing, Table S2). These proof-of-concept experiments demonstrate the potential utility of MN-QuIC as a portable, field-deployable diagnostic tool for researchers and agencies.

**Figure 5.**
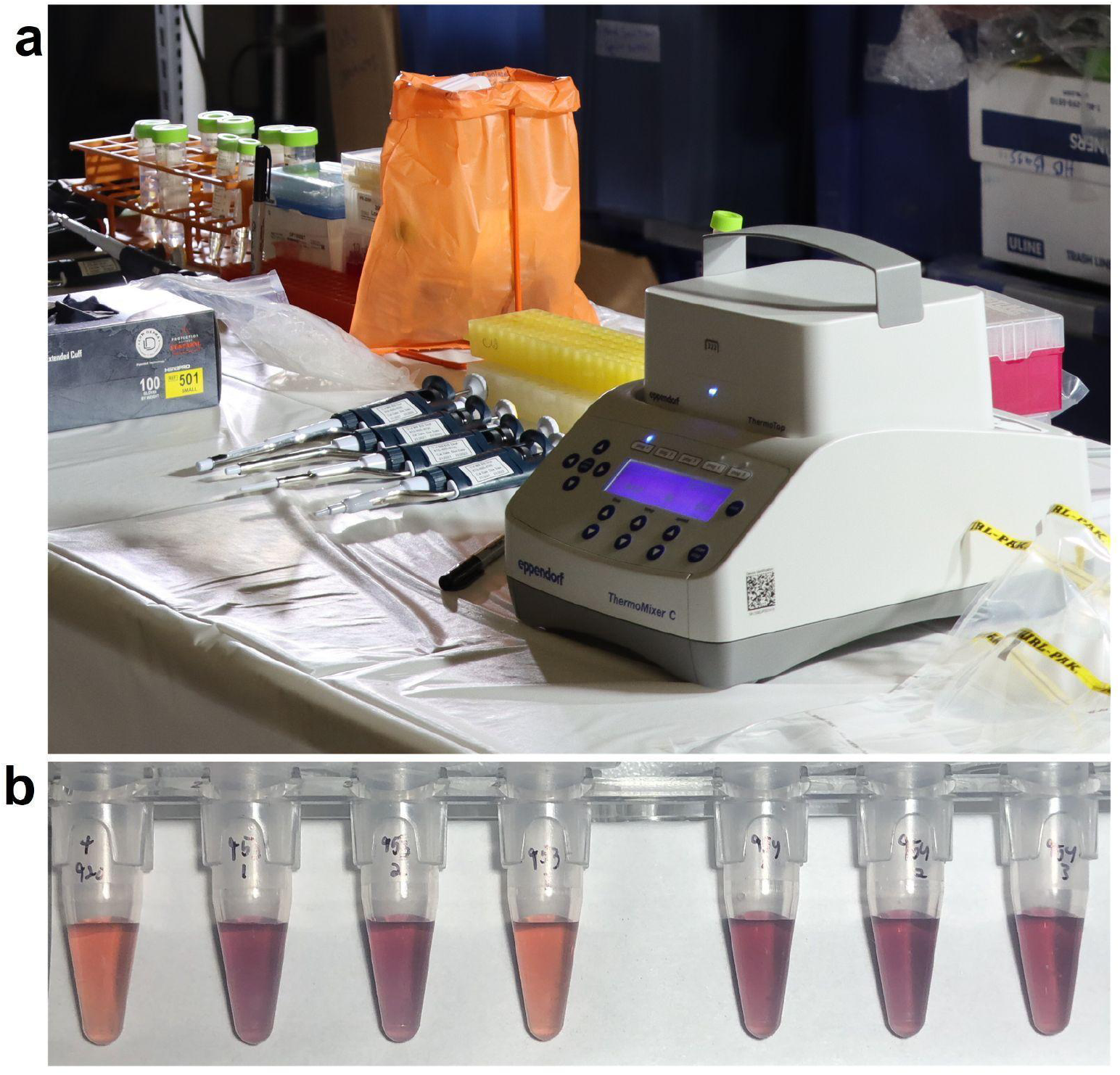
Images from field deployment. **a**. Image of portable MN-QuIC setup used during field deployment. **b**. Example of observed visible color of the MN-QuIC reaction when testing for CWD. Far-left tube (Tube 1) is a positive control, followed by CWD not-detected tube (in purple; Tubes 2, 3, 5, 6, 7) and a single CWD positive tube (in red; Tube 4). Image taken in the field.

In order to demonstrate higher throughput protocols, palatine tonsil samples from a set of ten CWD negative and ten CWD positive white-tailed deer (Table S3) were tested in a 96-well format. The status of these tissues was independently confirmed with RT-QuIC (Fig 6a). For MN-QuIC, each sample had eight replicates and was prepared and subjected to the QuIC protocol using a 96-well plate on a thermomixer for 24hrs. A multichannel pipette was then used to add the QuIC amplified protein to a separate 96-well plate filled with AuNP solution. Color changes were observed within the first minute. For RT-QuIC analyses, it is common practice to consider a particular sample positive if 50% or more of its wells are positive. ^24,51–53^ Using this approach, we successfully identified all 10 CWD-positive animals using MN-QuIC (Fig. 6b). CWD negative samples were identified using a threshold of <50% of wells being red (i.e., majority purple in color), and we correctly identified 100% of CWD negative samples with the MN-QuIC assay (Fig. 6b). In addition to visual color, these results were assessed by investigating the peak shift from the expected 517 nm (red) for positive samples and all red wells had absorbance peaks within 4nm of 517 nm.

**Figure 6:**
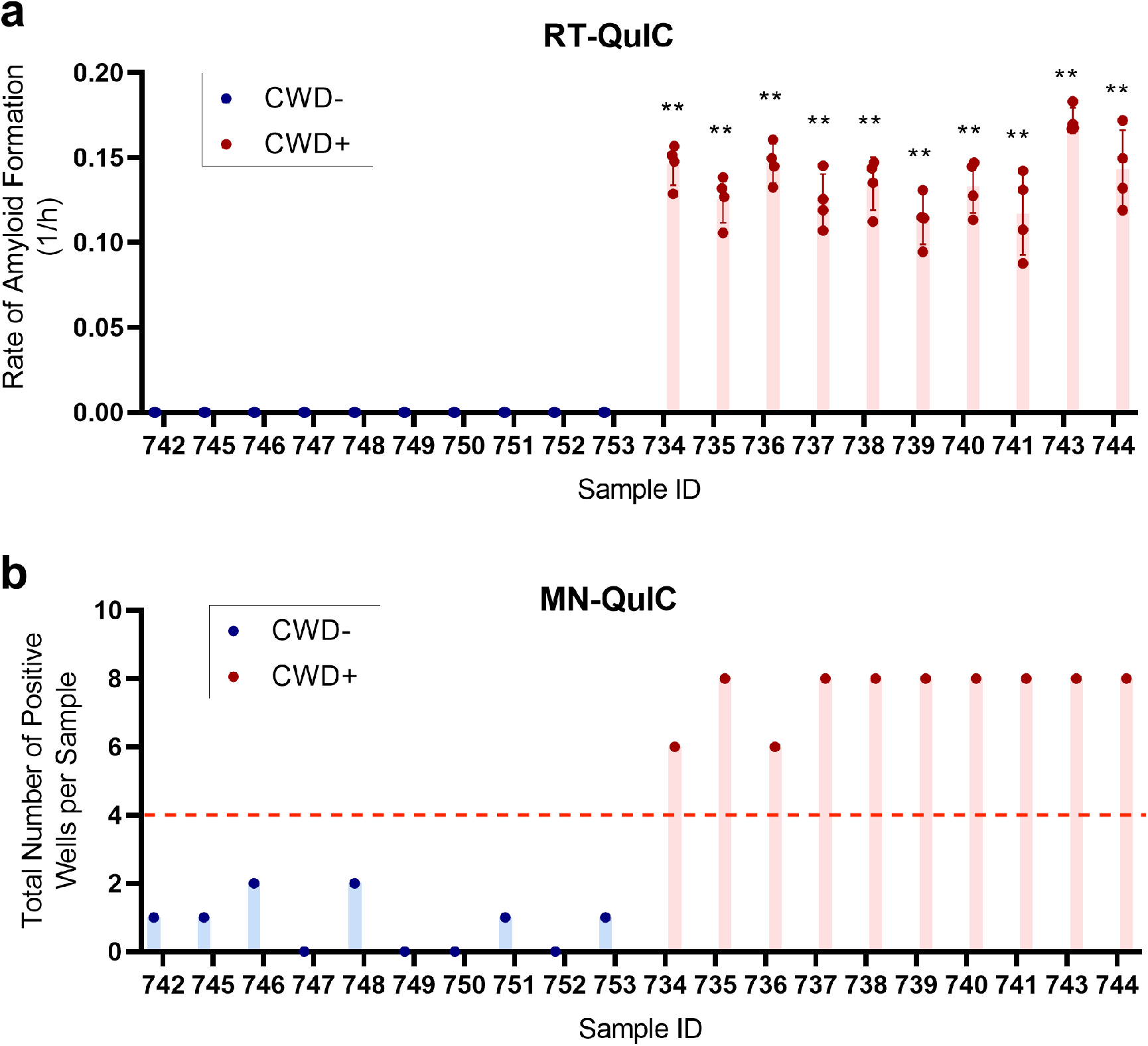
**a**. Fluorescent RT-QuIC data for tissues (see Supplementary Table 3) used in Panel b. **b**. MN-QuIC data for tonsil samples used in this study. Number of red wells out of the 8 replicates for each animal tested. Following published RT-QuIC protocols (see Results & Discussion) 50% or more of MN-QuIC red wells are identified as CWD positive and less than 50% of MN-QuIC red wells as CWD not detected or negative. **p<.01, error bars show standard deviation. Sample ID number on horizontal axis.

## Discussion

Given the continued spread of CWD among cervid populations throughout North America, Northern Europe, and South Korea ^13,14^ there is an urgent need to develop field-deployable diagnostic tools for CWD. Historically, AuNP colorimetric assays have been limited because of nonspecific binding and spontaneous aggregation issues when used with real-world samples.^43^ Here, we combined QuIC amplification of CWD prions with the simplicity of gold nanoparticles to eliminate nonspecific binding and spontaneous aggregation challenges and to enable the visualization of positive vs. negative CWD lymph node and palatine tonsil WTD samples. We hypothesized that the conformational differences between the native rPrP substrate and misfolded prion fibrils would influence the aggregation of AuNPs in solution.

By injecting post-QuIC-amplified protein solutions into AuNPs, we were able to clearly distinguish via visible color change between CWD positive and CWD negative medial retropharyngeal lymph node and palatine tonsil tissues. AuNP solutions for these samples appeared red and purple, respectively. We further confirmed that the color change was a result of the aggregation of gold nanoparticles by conducting DLS and TEM experiments that compared the effective particle sizes in the presence of native and misfolded rPrP. We observed that the AuNP aggregation was governed by electrostatic interactions by altering the pH of the solution. Finally, we demonstrated proof of concept experiments for the real-world utility of MN-QuIC by successfully identifying CWD-infected WTD tissues at a rural DNR field station in southeastern MN (Rushford, MN). In total, we examined 13 CWD positive and 24 CWD not-detected WTD that were independently tested using ELISA, IHC, and RT-QuIC technologies, and results secured with MN-QuIC were 100% consistent with these tests.

The primary laboratory equipment for MN-QuIC consists of a tissue homogenizer, temperature-controlled shaker, and if desired (but not necessary), a spectrometer for light absorbance readings in addition to visual observations. Compared to RT-QuIC and PMCA, which leverage ThT fluorescence and antibody-based Western blotting, respectively, ^6^ results from MN-QuIC can be visualized with the naked eye and quantified using straightforward light-absorbance readings. Because MN-QuIC is a protein amplification method, it can be adapted to use any tissue or biological sample that existing and future RT-QuIC protocols use, thus giving MN-QuIC wide versatility. Recent publications have used specially functionalized AuNPs to detect a variety of protein targets. ^39,54–56^ However, the AuNPs used here are capped with citrate, which is one of the most common methods for stabilization making it simple and widely commercially available. By eliminating the need for complicated/expensive visualization strategies used in RT-QuIC and PMCA, cervid managers, food processing plants, and smaller research labs have access to CWD diagnostics, an area previously accessible to only advanced research laboratories.

AuNPs have been used in a variety of advanced sensing applications. ^28,29,31,32,57–59^ Our work demonstrates that AuNPs can open promising avenues for the identification of misfolded prions. Because prion proteins have strong interactions with simply functionalized metallic surfaces, besides AuNPs, we envision a broad range of metallic nanoparticles with various materials and shapes to be useful in detection. Additionally, substrate-based nanostructures exhibiting optical resonances could be useful in detecting conformational changes via other sensing modalities such as surface-enhanced infrared absorption spectroscopy ^60,61^ to further improve the speed and accuracy of prion detection. Examining other areas of the electromagnetic spectrum, such as tera-Hertz sensing, could also lead to improved detection. ^62^

To demonstrate field deployment capabilities of the MN-QuIC assay, we collected medial retropharyngeal lymph nodes, parotid lymph nodes, and palatine tonsil samples from 13 WTD at a rural DNR field station. Using both pooled and individual tissues of these 13 individuals, MN-QuIC was 100% successful in identifying three CWD positive and 10 CWD not-detected animals. The successful field-based classification of these animals provides clear proof of concept demonstration of MN-QuICs utility as a portable, sensitive field test. We note that our MN-QuIC analyses have yet to produce statistically significant false-negative results. This observation is critically important when considering MN-QuIC as a field-based diagnostic tool for CWD. Moreover, any positive result can be independently validated using downstream RT-QuIC, ELISA, and/or IHC testing. Given growing concerns of CWD prion strain variation and risks to human and animal health, ^15^ any field-based diagnostic assay that avoids producing false-negative is preferred.

RPLN and palatine tonsils collected from WTD were the basis for the laboratory-based analyses conducted herein because these tissues are ideal for early and accurate identification of CWD infection, with tonsils additionally having antemortem applications. ^27,63^ Future analyses will focus on leveraging MN-QuIC for CWD diagnostics using a variety of antemortem biological samples (e.g., blood, saliva, and feces). RT-QuIC amplification assays using samples acquired from living deer have recently been reported ^45,64^ and these assays could readily be combined with MN-QuIC to provide field-deployable antemortem tests of both wild and farmed cervids. Moreover, MN-QuIC may have potential food-safety test applications given the recent documentation of RT-QuIC-based detection of CWD prions in WTD muscles used for human and animal consumption. ^26^

Despite the critical importance of prion disease, the mechanisms underlying the interactions between AuNPs and prions are relatively underexplored to date, likely due to the difficulty of handling potentially infectious samples. Detailed characterization of their interactions will be highly desirable to further improve the utility of these assays. Based on the available knowledge in the field, in this study, we hypothesized that as the pH approaches the prion’s isoelectric point, the electrostatic force of attraction between the negatively charged citrate capped AuNP and the protein would decrease. Our analyses revealed that as the pH neared the theoretical isoelectric point of rPrP, the wavelength of the peak absorbance of the AuNPs spiked with native (CWD negative) protein decreased. We also found that pH alterations had little effect on AuNP solutions without protein, indicating that the effect of pH on native rPrP-spiked AuNP solutions was not caused by intrinsic AuNP aggregation in response to the changing pH. QuIC-amplified CWD positive solutions did not change with varying pH because of fibril formation effects on free-floating prion concentration. Therefore, the difference in interactions between CWD negative and positive solutions is likely governed by electrostatic forces and rPrP concentration effects. However, other factors such as hydrophobic interactions ^65,66^ could also play a role. Various studies have shown that both native and misfolded PrP bind various metal ions and bulk metals including gold. ^16,40–42^ Our research reveals AuNPs stabilized with a simple citrate capping readily interact with the truncated rPrP substrate that is used as the primary substrate for a growing variety of QuIC assays. Future structural analyses focused on native rPrP and prion fibrils ^67^ could provide further insight into how native rPrP, but not misfolded rPrP fibrils, influence AuNP aggregation.

## Conclusions

Our assay holds great promise not only for the visual detection of CWD-positive samples but also for the detection of other protein misfolding diseases. The need for inexpensive, sensitive, widely deployable diagnostics for neurodegenerative diseases is only growing as neurodegenerative diseases are predicted to greatly increase in the next decades. ^2,68^ It has been proposed that advances in CWD diagnostics will yield technologies that are useful for a broad range of neurodegenerative diseases. ^6^ RT-QuIC protocols have already been developed for a number of sample types allowing for antemortem tests.^45,64^ These and future amplification methods could readily be combined with MN-QuIC. Additionally, QuIC amplification protocols have been developed for a variety of other protein misfolding diseases including scrapie in sheep^69^, BSE in cattle^70^, and Alzheimer’s^71^, Parkinson’s^52,72^, and CJD in humans.^22,23^ Thus, we posit that the combination of AuNP technology with protein amplification assays has great potential for the development of versatile neurodegenerative disease diagnostic platforms. By eliminating the need for expensive/complicated visualization schemes, our hybrid assay technology has the potential to greatly increase access to neurodegenerative disease diagnostics. It is our vision that in the future, variations of this AuNP-based protein amplification/detection assay could be deployed in medical clinics around the world to aid in neurodegenerative diagnosis and early application of therapeutics.

## Methods

### Tissue preparation

Twenty-eight WTD tissues (14 CWD-negative and 14 CWD-positive) were selected for laboratory-based RT-QuIC and MN-QuIC analyses. These samples were collected from WTD through collaboration with the Minnesota DNR (Schwabenlander *et al*.^27^ 2021; Tables S1 and S3), and their CWD status was independently identified utilizing the Bio-Rad TeSeE Short Assay Protocol (SAP) Combo Kit (BioRad Laboratories Inc., Hercules, CA, USA). Positive RPLNs were confirmed by IHC at the Colorado State University Veterinary Diagnostic Laboratory). Metadata containing information of all specimens examined in the lab, including tissue type, is provided in supplementary materials (Tables S1 and S3). WTD RPLNs and palatine tonsils were homogenized in PBS (10% w:v) in 2 mL tubes containing 1.5 mm zirconium beads with a BeadBug Homogenizer (Benchmark Scientific, Sayreville New Jersey, USA) on max speed for 90 sec. These samples are referred to as 10% homogenates. All CWD positive and negative samples were selected based on independent ELISA, IHC, and/or RT-QuIC results and were subsampled using methods as reported in Schwabenlander *et al*. ^27^

### Preparation of recombinant substrate

Recombinant hamster PrP (HaPrP90-231) production and purification followed the methods in Schwabenlander *et al*. ^27^ The substrate is derived from a truncated form (amino acids 90-231) of the Syrian hamster PRNP gene cloned into the pET41-a(+) expression vector and was expressed in Rosetta (DE3) *E. coli*. The original clone was provided by the National Institutes of Health Rocky Mountain Laboratory.

### RT-QuIC for lymph tissues and spontaneous misfolding of rPrP

For QuIC analysis, a master mix was made to the following specifications: 1X PBS, 1mM Ethylenediaminetetraacetic acid (EDTA), 170mM NaCl, 10 μM thioflavin T (ThT), and 0.1 mg/mL rPrP. In instances where the end reaction would be analyzed using AuNPs, ThT could be excluded. The 10% tissue homogenates (prepared as described above) were further diluted 100-fold in 0.1% Sodium Dodecyl Sulfate (SDS) using methods from Schwabenlander *et al*. ^27^ (final tissue dilution: 0.1%), and 2 μL of the diluent were added to each well containing 98 uL of master mix. Spontaneous misfolding of recombinant prion protein was generated similarly but with unfiltered recombinant proteins and reagents. For these reactions, no infectious seed was necessary. The spontaneously misfolded material was used to seed reactions for the dynamic light scattering experiment, described below. Plates were amplified on a FLUOstar® Omega plate reader (BMG Labtech, Cary, North Carolina, USA; 42°C, 700 rpm, double orbital, shake for 57 s, rest for 83 s). Fluorescent readings were taken at ∼45 min increments.

### Thermomixer-based amplification

We leveraged a standard benchtop shaking incubator (thermomixer) to produce QuIC-based prion amplifications as previously reported by Cheng *et al*. ^22^ and Vendramelli *et al*, ^23^ although with slight modifications. Plates were prepared identically to those amplified on the plate reader (see protocol above). Instead of shaking on a plate reader, reactions were performed on a ThermoMixer® C equipped with SmartBlock plate and Thermotop (Eppendorf, Enfield, Connecticut, USA) at 48°C for 24hrs at 600 RPM (60s shake and 60s rest). We selected a 24 hour run time based on independent RT-QuIC results for RPLNs and palatine tonsils from CWD positive WTD reported in Schwabenlander *et al*. ^27^, including those examined herein, showing significant seeding activity within 9 to 24 hours (Fig. S3). The resultant products were visualized with the addition of gold nanoparticles (as described below).

### Preparation of gold nanoparticles

Post-amplified material was visualized with 15 nm citrate-capped gold nanoparticles purchased from Nanopartz (Loveland, Colorado, USA) with stock concentrations ranging from 2.45nM to 2.7nM. AuNP protocols were modified from Špringer *et al*. ^73^ and Zhang *et al*. ^39^ AuNPs were buffer exchanged using 530ul of stock solution that was centrifuged in 1.6mL tubes at 13,800g for 10 min. 490ul of supernatant was removed and the undisturbed pellet was resuspended with 320ul of a low concentration phosphate buffer (PBS_low_; pH 7.4 via addition of HCl) made of 10mM Na_2_HPO_4_(Anhydrous), 2.7mM KCl, 1.8mM KH_2_PO_4_). After the quaking/incubation steps, protein solutions were diluted to 50% in MN-QuIC buffer (pH 7.2), consisting of 1X PBS with the addition of final concentrations of 1mM EDTA, 170mM NaCl, 1.266mM sodium phosphate. Forty microliters of the protein diluted 50% in MN-QuIC buffer were then added to the 360ul AuNP solution with ample mixing (results shown in Fig 3b,c). This solution was left to react at room temperature (RT) for 30 min (although a visible color change is observable within 60 sec) before visual color was recorded (purple or red) and photographed. After images were taken, three replicates of 100ul were taken from the 400ul AuNP mixture and pipetted into three separate wells of a 96-well plate. The absorbance spectrum was then recorded for each well at wavelengths 400-800nm using the FLUOstar® Omega plate reader (BMG Labtech, Cary, North Carolina, USA). For AuNP visualization experiments performed to determine higher throughput capacity (Fig 6), proteins were prepared in the same way. After amplification on the thermomixer, proteins on a 96-well plate were diluted to 50% using MN-QuIC buffer and a multichannel pipette. 90ul of AuNPs were then added to a separate non-binding 96-well plate. The AuNP wells were spiked with 10ul of the diluted protein from the thermomixer (post-amplification) using a multichannel pipette. After waiting 30min, the color changes were observed and the absorbance spectrum of the plate was taken.

### Dynamic light scattering

Spontaneously misfolded rPrP samples (described above) were produced from solutions of rPrP with no seed added. In addition to these samples, a 96-well RT-QuIC reaction was performed with half the wells consisting of native rPrP seeded with spontaneously misfolded rPrP, and half consisting of native rPrP with no seed. The 96-well plate was then amplified using QuIC protocols described above. Post-amplification, seeded samples were confirmed to have fibrillation while the non-seeded samples were confirmed to not have fibrillation based on ThT binding (described above). Seeded and non-seeded samples were diluted to 50% in MN-QuIC buffer, and 40ul of these solutions were added to 360ul of AuNPs in PBS_low_. Additionally, a blank with no protein was produced by adding 40ul of MN-QuIC buffer to 360ul of AuNPs in PBS_low_. For native rPrP samples, color change was observed within 1 min of rPrPaddition. No color change was observed in spontaneously misfolded rPrP samples at any time length. Dynamic light scattering measurements of all samples were taken after 5 min of protein addition using a Microtract NanoFlex Dynamic Light Scattering Particle Analyzer (Verder Scientific, Montgomeryville, PA, USA), and measurement times were 60 seconds. Five measurements were taken for each sample and then averaged.

### TEM measurements

TEM measurements were performed using a Tecnai G2 F30 Transmission Electron Microscope. Images were recorded on a Gatan K2 Summit direct electron detector. 20ul of post-amplified positive and negative rPrP were added to separate volumes of 200ul of AuNPs prepared as described above. Each protein solution was added to separate uncharged TEM grids. Stain was not used because it was unnecessary to view the AuNPs.

### Effects of pH and rPrP substrate concentration on the AuNP-protein interaction

In order to test the effects of pH on the interaction of rPrP with AuNP, four different 10mM tris-buffer solutions with pHs ranging from 7.2 to 9.0 were created. Tris was used to give buffering for the desired pH range. AuNPs were buffer exchanged as described above except tris-buffer was used instead of PBS_low_. Protein solutions were added as previously described. It can be noted that the peak absorbance for the tris buffer solution below pH 7.4 is still not as high as the peak shifts in pH 7.4 PBS_low_. This is likely due to the differences in tris and PBS_low_ buffers.

To examine effects of rPrP substrate concentration (Fig. S2), 6 different master mixes were made with concentrations of native rPrP ranging from 0mg/ml to 0.1mg/ml. 10ul of each solution were added to separate wells containing 90ul AuNPs (pH 7.4 AuNPs prepared as described above).

### Additional statistical information

GraphPad Prism version 9.0 for Windows (GraphPad Software, San Diego, California USA, www.graphpad.com) was used for conducting statistical analysis. Three technical replicates were used to demonstrate the potential application of AuNP on spontaneously misfolded rPrP. For initial trials on RPLN tissues from eight (four positive and four negative) animals, four and three technical replicates were used for RT-QuIC and AuNPs, respectively. For plate-based protocols, we tested palatine tonsils from ten positive and ten negative animals using four and eight replicates for RT-QuIC and AuNPs, respectively. Unless specified in figures, rate of amyloid formation and maximum wavelength of samples were compared to negative controls on their respective plate. The one-tailed Mann-Whitney unpaired u-test (α=0.05) was used to test the average difference for all parameters of interests between samples.

### Field deployment

In March of 2021, we collaborated with the Minnesota DNR during annual CWD surveillance of the wild WTD population in Fillmore and Winona Counties, Minnesota. We assembled the necessary MN-QuIC equipment as described above on two portable tables within a DNR facility in Rushford, MN. RPLNs, parotid lymph nodes, and palatine tonsil were collected as described in Schwabenlander *et al*.,^27^ and were sampled and pooled together for each of the 13 animals tested. Tissues for suspected positive animals were tested individually (Table S2). All tissues were subject to 24hr MN-QuIC protocols as described above. Three replicates were performed for each of the 13 animals and, for field-based analyses, an animal was considered CWD positive if one or more replicates was red. An animal was considered CWD not-detected if all three replicates were blue or purple.

## ASSOCIATED CONTENT

### Supporting Information

Specimens examined, details on RT-QuIC and MN-QuIC assay results, and prion seeding activity (PDF)

## Author Contributions

PRC and ML conceived the study. PRC, ML, and GR performed molecular experiments. PRC, ML, MS, SHO, and PAL assisted with experimental design and interpreted the results. PRC, ML, MS, TMW, and PAL conducted field-based testing of the diagnostics presented herein. PRC and ML performed statistical analyses. SHO and PAL oversaw the research. All authors wrote and contributed to the final manuscript.

## Acknowledgments

We thank Christopher Ertsgaard and Dong Jun Lee for helpful discussions on experimental results and protocols. Portions of this work were conducted in the Minnesota Nano Center, which is supported by the National Science Foundation through the National Nano Coordinated Infrastructure Network (NNCI) under Award Number ECCS-2025124. W. Zhang provided the expertise for TEM studies. These studies were carried out in the University of Minnesota Characterization Facility, which receives partial support from the NSF through the MRSEC (Award Number DMR-2011401) and the NNCI (Award Number ECCS-2025124) programs. F. Schendel, T. Douville, and staff of the University of Minnesota Biotechnology Resource Center provided critical support concerning the large-scale production of recombinant proteins. We thank the Minnesota Department of Natural Resources, especially M. Carstensen, Lou Cornicelli, E. Hildebrand, P. Hagen, and K. LaSharr, for providing the white-tailed deer tissues used for our analyses and logistical assistance for MN-QuIC field deployment. K. Wilson of the Colorado State University Veterinary Diagnostic Laboratory provided assistance with ELISA and IHC testing of samples reported herein. S. Stone provided valuable logistical assistance with our molecular work. We thank NIH Rocky Mountain Labs, especially B. Caughey, A. Hughson, and C. Orru for training and assistance with the implementation of RT-QuIC and for supplying the rPrP clone. Funding for research performed herein was provided by the Minnesota Department of Natural Resources, the Minnesota State Legislature through the Minnesota Legislative-Citizen Commission on Minnesota Resources (LCCMR), Minnesota Agricultural Experiment Station Rapid Agricultural Response Fund, and start-up funds awarded to P.A.L. through the Minnesota Agricultural, Research, Education, Extension and Technology Transfer (AGREETT) program. Fig. 1 and parts of Fig. 2 were created using BioRender (BioRender.com).

## Notes

The authors declare no competing financial interest.

## References

1. Prusiner, S. B. Nobel Lecture: Prions. Proc. Natl. Acad. Sci. vol. 95 13363–13383 (1998).

2. Telling, G. C. Breakthroughs in antemortem diagnosis of neurodegenerative diseases. Proc. Natl. Acad. Sci. vol. 116 22894–22896 (2019).

3. Collinge, J. Prion diseases of humans and animals: their causes and molecular basis. Annu. Rev. Neurosci. 24, 519–550 (2001).

4. Williams, E. S. & Young, S. Chronic wasting disease of captive mule deer: a spongiform encephalopathy. J. Wildl. Dis. 16, 89–98 (1980).

5. Fortin, J. S. et al. Equine pituitary pars intermedia dysfunction: a spontaneous model of synucleinopathy. Sci. Rep. 11, 16036 (2021).

6. Haley, N. J. & Richt, J. A. Evolution of Diagnostic Tests for Chronic Wasting Disease, a Naturally Occurring Prion Disease of Cervids. Pathogens 6, (2017).

7. Haley, N. J. et al. Chronic wasting disease management in ranched elk using rectal biopsy testing. Prion 12, 93–108 (2018).

8. Martinez, B. & Peplow, P. V. MicroRNAs in Parkinson’s disease and emerging therapeutic targets. Neural Regeneration Res. 12, 1945–1959 (2017).

9. Hajipour, M. J. et al. Advances in Alzheimer’s Diagnosis and Therapy: The Implications of Nanotechnology. Trends Biotechnol. 35, 937–953 (2017).

10. Figgie, M. P., Jr & Appleby, B. S. Clinical Use of Improved Diagnostic Testing for Detection of Prion Disease. Viruses 13, (2021).

11. Parnetti, L. et al. CSF and blood biomarkers for Parkinson’s disease. Lancet Neurol. 18, 573–586 (2019).

12. Prusiner, S. B. et al. Scrapie Prions Aggregate to Form Amyloid-Like Birefringent Rods. Cell 35, 349–358 (1983).

13. Hannaoui, S., Schatzl, H. M. & Gilch, S. Chronic wasting disease: Emerging prions and their potential risk. PLoS Pathog. 13, e1006619 (2017).

14. Joly, D. O. et al. Chronic wasting disease in free-ranging Wisconsin White-tailed Deer. Emerg. Infect. Dis. 9, 599–601 (2003).

15. Osterholm, M. T. et al. Chronic Wasting Disease in Cervids: Implications for Prion Transmission to Humans and Other Animal Species. MBio 10, e01091–19 (2019).

16. Westergard, L., Christensen, H. M. & Harris, D. A. The cellular prion protein (PrPC): Its physiological function and role in disease. Biochim Biophys Acta 1772, 629–644 (2007).

17. U.S. Fish and Wildlife Service & U.S. Census Bureau. 2016 National Survey of Fishing, Hunting and Wildlife-Associated Recreation. https://www.census.gov/library/publications/2018/demo/fhw-16-nat.html. (2018).

18. Wu, F. et al. Deer antler base as a traditional Chinese medicine: a review of its traditional uses, chemistry and pharmacology. J. Ethnopharmacol. 145, 403–415 (2013).

19. Safar, J. G. et al. Prion clearance in bigenic mice. J. Gen. Virol. 86, 2913–2923 (2005).

20. Haley, N. J. et al. Sensitivity of protein misfolding cyclic amplification versus immunohistochemistry in ante-mortem detection of chronic wasting disease. J. Gen. Virol. 93, 1141–1150 (2012).

21. Saborio, G. P., Permanne, B. & Soto, C. Sensitive detection of pathological prion protein by cyclic amplification of protein misfolding. Nature 411, 810–813 (2001).

22. Cheng, K. et al. Endpoint Quaking-Induced Conversion: a Sensitive, Specific, and High-Throughput Method for Antemortem Diagnosis of Creutzfeldt-Jacob Disease. J. Clin. Microbiol. 54, 1751–1754 (2016).

23. Vendramelli, R., Sloan, A., Simon, S. L. R., Godal, D. & Cheng, K. ThermoMixer-Aided Endpoint Quaking-Induced Conversion (EP-QuIC) Permits Faster Sporadic Creutzfeldt-Jakob Disease (sCJD) Identification than Real-Time Quaking-Induced Conversion (RT-QuIC). J. Clin. Microbiol. 56, e00423–18 (2018).

24. Cheng, Y. C. et al. Early and Non-Invasive Detection of Chronic Wasting Disease Prions in Elk Feces by Real-Time Quaking Induced Conversion. PLoS One 11, e0166187 (2016).

25. Atarashi, R. et al. Ultrasensitive human prion detection in cerebrospinal fluid by real-time quaking-induced conversion. Nat. Med. 17, 175–178 (2011).

26. Li, M. et al. RT-QuIC detection of CWD prion seeding activity in white-tailed deer muscle tissues. Sci. Rep. 11, 16759 (2021).

27. Schwabenlander, M. D. et al. Comparison of Chronic Wasting Disease Detection Methods and Procedures: Implications for Free-Ranging White-Tailed Deer (Odocoileus Virginianus) Surveillance and Management. J. Wildl. Dis. (2021) doi:10.7589/JWD-D-21-00033.

28. Tsai, T.-T. et al. Diagnosis of Tuberculosis Using Colorimetric Gold Nanoparticles on a Paper-Based Analytical Device. ACS Sens 2, 1345–1354 (2017).

29. Pelaz, B. et al. Diverse Applications of Nanomedicine. ACS Nano 11, 2313–2381 (2017).

30. Howes, P. D., Chandrawati, R. & Stevens, M. M. Bionanotechnology. Colloidal nanoparticles as advanced biological sensors. Science 346, 1247390 (2014).

31. Thiramanas, R. & Laocharoensuk, R. Competitive binding of polyethyleneimine-coated gold nanoparticles to enzymes and bacteria: a key mechanism for low-level colorimetric detection of gram-positive and gram-negative bacteria. Microchim. Acta 183, 389–396 (2016).

32. Du, X.-J., Zhou, T.-J., Li, P. & Wang, S. A rapid Salmonella detection method involving thermophilic helicase-dependent amplification and a lateral flow assay. Mol. Cell. Probes 34, 37–44 (2017).

33. Mayer, K. M. & Hafner, J. H. Localized surface plasmon resonance sensors. Chem. Rev. 111, 3828–3857 (2011).

34. Myroshnychenko, V. et al. Modelling the optical response of gold nanoparticles. Chem. Soc. Rev. 37, 1792–1805 (2008).

35. Lal, S., Link, S. & Halas, N. J. Nano-optics from sensing to waveguiding. Nature Photonics vol. 1 641–648 (2007).

36. Sepúlveda, B., Angelomé, P. C., Lechuga, L. M. & Liz-Marzán, L. M. LSPR-based nanobiosensors. Nano Today 4, 244–251 (2009).

37. Dahlin, A. et al. Localized surface plasmon resonance sensing of lipid-membrane-mediated biorecognition events. J. Am. Chem. Soc. 127, 5043–5048 (2005).

38. Zhao, W., Brook, M. A. & Li, Y. Design of gold nanoparticle-based colorimetric biosensing assays. ChemBioChem 9, 2363–2371 (2008).

39. Zhang, H.-J. et al. Gold nanoparticles as a label-free probe for the detection of amyloidogenic protein. Talanta 89, 401–406 (2012).

40. Flechsig, E. et al. Transmission of scrapie by steel-surface-bound prions. Molecular Medicine 7, 679–684 (2001).

41. Weissmann, C., Enari, M., Klöhn, P. C., Rossi, D. & Flechsig, E. Transmission of prions. J. Infec. Dis. 186, S157–S165 (2002).

42. Luhr, K. M., Löw, P., Taraboulos, A., Bergman, T. & Kristensson, K. Prion adsorption to stainless steel is promoted by nickel and molybdenum. J. Gen. Virol. 90, 2821–2828 (2009).

43. Masson, J.-F. Surface Plasmon Resonance Clinical Biosensors for Medical Diagnostics. ACS Sens 2, 16–30 (2017).

44. Henderson, D. M. et al. Quantitative assessment of prion infectivity in tissues and body fluids by real-time quaking-induced conversion. J. Gen. Virol. 96, 210–219 (2015).

45. Tennant, J. M. et al. Shedding and stability of CWD prion seeding activity in cervid feces. PLoS One 15, e0227094 (2020).

46. Henderson, D. M. et al. Progression of chronic wasting disease in white-tailed deer analyzed by serial biopsy RT-QuIC and immunohistochemistry. PLoS One 15, e0228327 (2020).

47. Wang, A., Perera, Y. R., Davidson, M. B. & Fitzkee, N. C. Electrostatic Interactions and Protein Competition Reveal a Dynamic Surface in Gold Nanoparticle–Protein Adsorption. J. Phys. Chem. C 120, 24231–24239 (2016).

48. Anika Gladytz Bernd Abel & Risselada, H. J. Gold-induced fibril growth: the mechanism of surface-facilitated amyloid aggregation. Angew. Chem. Int. Ed. 55, 11242–11246 (2016).

49. Gasteiger, E. et al. Protein Identification and Analysis Tools on the ExPASy Server. The Proteomics Protocols Handbook 571–607 (2005) doi:10.1385/1-59259-890-0:571.

50. Csapó, E. et al. Surface and Structural Properties of Gold Nanoparticles and Their Biofunctionalized Derivatives in Aqueous Electrolytes Solution. J. Dispers. Sci. Technol. 35, 815–825 (2014).

51. Haley, N. J., Henderson, D. M., Senior, K., Miller, M. & Donner, R. Evaluation of Winter Ticks (Dermacentor albipictus) Collected from North American Elk (Cervus canadensis) in an Area of Chronic Wasting Disease Endemicity for Evidence of PrPCWD Amplification Using Real-Time Quaking-Induced Conversion Assay. mSphere 6, e0051521 (2021).

52. Rossi, M. et al. Ultrasensitive RT-QuIC assay with high sensitivity and specificity for Lewy body-associated synucleinopathies. Acta Neuropathol. 140, 49–62 (2020).

53. Mok, T. H. et al. Bank vole prion protein extends the use of RT-QuIC assays to detect prions in a range of inherited prion diseases. Sci. Rep. 11, 5231 (2021).

54. Zhang, X. Gold Nanoparticles: Recent Advances in the Biomedical Applications. Cell Biochem. Biophys. 72, 771–775 (2015).

55. Li, J., Yan, X., Li, X., Zhang, X. & Chen, J. A new electrochemical immunosensor for sensitive detection of prion based on Prussian blue analogue. Talanta 179, 726–733 (2018).

56. Zhao, J. et al. Graphene oxide-gold nanoparticle-aptamer complexed probe for detecting amyloid beta oligomer by ELISA-based immunoassay. J. Immunol. Methods 489, 112942 (2021).

57. Ruppert, C., Phogat, N., Laufer, S., Kohl, M. & Deigner, H.-P. A smartphone readout system for gold nanoparticle-based lateral flow assays: application to monitoring of digoxigenin. Microchim. Acta 186, 119 (2019).

58. Mahato, K. et al. Gold nanoparticle surface engineering strategies and their applications in biomedicine and diagnostics. 3 Biotech 9, 57 (2019).

59. Lee, J.-H., Cho, H.-Y., Choi, H. K., Lee, J.-Y. & Choi, J.-W. Application of Gold Nanoparticle to Plasmonic Biosensors. Int. J. Mol. Sci. 19, (2018).

60. Adato, R. et al. Ultra-sensitive vibrational spectroscopy of protein monolayers with plasmonic nanoantenna arrays. Proc. Natl. Acad. Sci. U. S. A. 106, 19227–19232 (2009).

61. Altug, H., Oh, S.-H., Maier, S. A. & Homola, J. Advances and applications of nanophotonic biosensors. Nat. Nanotechnol. 17, 5–16 (2022).

62. Heo, C. et al. Identifying Fibrillization State of Aβ Protein via Near-Field THz Conductance Measurement. ACS Nano 14, 6548–6558 (2020).

63. Wild, M. A., Spraker, T. R., Sigurdson, C. J., O’Rourke, K. I. & Miller, M. W. Preclinical diagnosis of chronic wasting disease in captive mule deer (Odocoileus hemionus) and white-tailed deer (Odocoileus virginianus) using tonsillar biopsy. J. Gen. Virol. 83, 2629–2634 (2002).

64. Ferreira, N. C. et al. Detection of chronic wasting disease in mule and white-tailed deer by RT-QuIC analysis of outer ear. Sci. Rep. 11, 7702 (2021).

65. Gao, D. et al. Studies on the interaction of colloidal gold and serum albumins by spectral methods. Spectrochim. Acta A Mol. Biomol. Spectrosc. 62, 1203–1208 (2005).

66. Szekeres, G. P. & Kneipp, J. Different binding sites of serum albumins in the protein corona of gold nanoparticles. Analyst 143, 6061–6068 (2018).

67. Kraus, A. et al. High-resolution structure and strain comparison of infectious mammalian prions. Mol. Cell 81, 4540–4551.e6 (2021).

68. Matthews, K. A. et al. Racial and ethnic estimates of Alzheimer’s disease and related dementias in the United States (2015-2060) in adults aged ≥65 years. Alzheimers. Dement. 15, 17–24 (2019).

69. Orrú, C. D. et al. Human variant Creutzfeldt-Jakob disease and sheep scrapie PrP(res) detection using seeded conversion of recombinant prion protein. Protein Eng. Des. Sel. 22, 515–521 (2009).

70. Hwang, S., Greenlee, J. J. & Nicholson, E. M. Use of bovine recombinant prion protein and real-time quaking-induced conversion to detect cattle transmissible mink encephalopathy prions and discriminate classical and atypical L-and H-Type bovine spongiform encephalopathy. PLoS One 12, e0172391 (2017).

71. Kraus, A. et al. Seeding selectivity and ultrasensitive detection of tau aggregate conformers of Alzheimer disease. Acta Neuropathol. 137, 585–598 (2019).

72. Fairfoul, G. et al. Alpha-synuclein RT-QuIC in the CSF of patients with alpha-synucleinopathies. Ann Clin Transl Neurol 3, 812–818 (2016).

73. Špringer, T. & Homola, J. Biofunctionalized gold nanoparticles for SPR-biosensor-based detection of CEA in blood plasma. Anal. Bioanal. Chem. 404, 2869–2875 (2012).

