## Supplementary Information for "A Field-Deployable Diagnostic Assay for the Visual Detection of Misfolded Prions"

### Supplemental Results

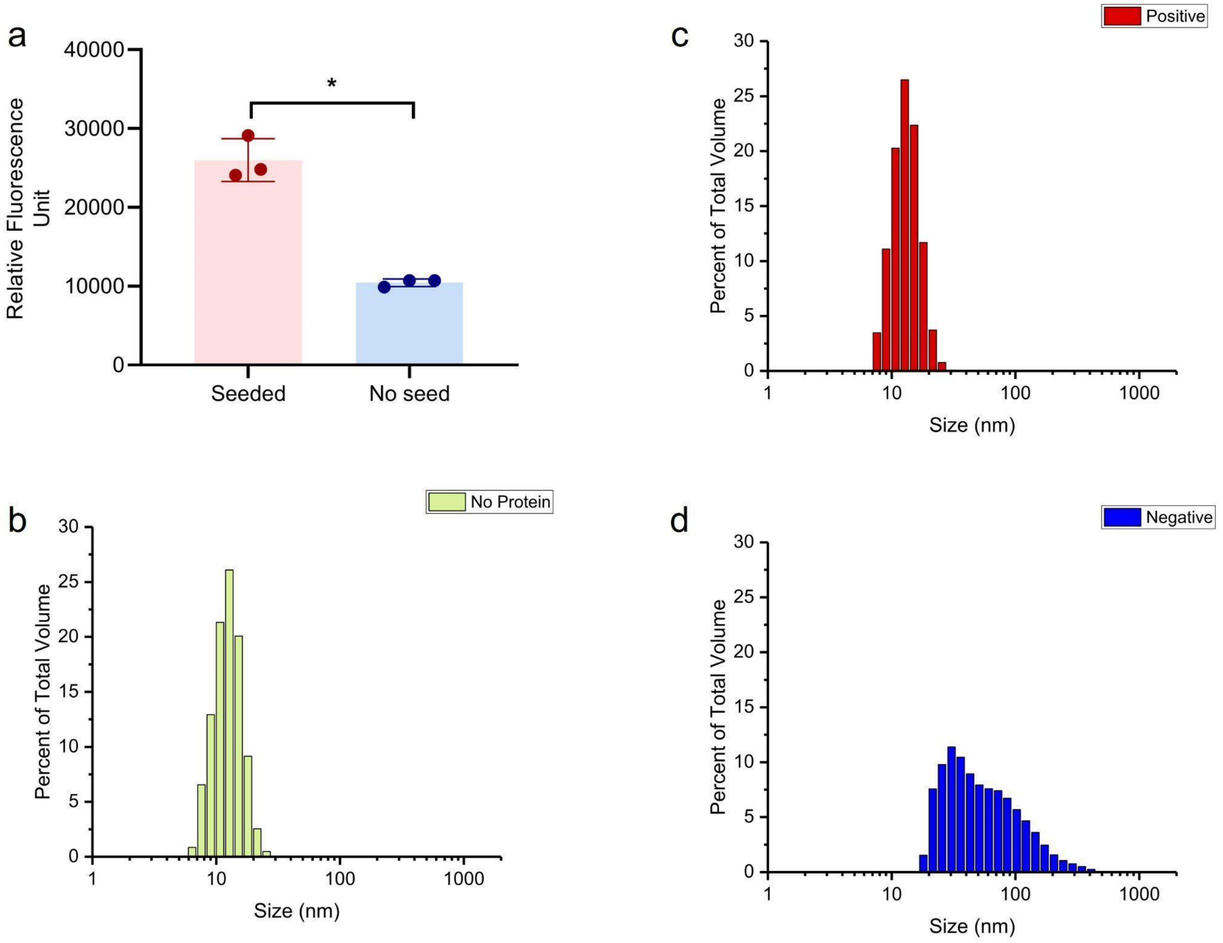


**Supplementary Figure 1.** ThT fluorescence, light absorbance, and average particle size measurements for native and misfolded recombinant hamster prion protein. **a.** The relative fluorescence units (ThT fluorescence) of post QuIC solutions containing misfolded protein seeds and solutions without misfolded protein seeds. **b.** Distribution of DLS readings of AuNP solution with no protein added. Reported as percent of total volume of AuNPs present. **c.** Distribution of DLS readings of AuNP solution with misfolded (positive) rHaPrP. **d.** Distribution of DLS readings of AuNP solution with native (negative) rHaPrP. *, $p$-value $< 0.05$, error bars show standard deviation.

**Supplementary Table 1. Metadata of Examined Wild White-Tailed Deer RPLN.** MNPRO = Minnesota Center for Prion Research and Outreach. RPLN = medial retropharyngeal lymph node. Official CWD test result indicates the RPLN was independently examined using ELISA and/or IHC testing at Colorado State University (see Schwabenlander et al. 2021, citation 26 in main text). ND=Not Detected.

| Animal MNPRO ID | Age | Sex | Official CWD Test Result | RT-QuIC Result | MN-QuIC Results |
|---|---|---|---|---|---|
| 166 | Adult | Female | Positive | Positive | Positive |
| 250 | Adult | Male | Positive | Positive | Positive |
| 360 | Adult | Female | Positive | Positive | Positive |
| 508 | Adult | Male | Positive | Positive | Positive |
| 922 | Adult | Female | ND | ND | ND |
| 924 | Adult | Female | ND | ND | ND |
| 925 | Fawn | Female | ND | ND | ND |
| 926 | Fawn | Female | ND | ND | ND |

**Supplementary Table 2. Metadata of Examined Wild White-Tailed Deer in Field Test.** RPLN=medial retropharyngeal lymph node. Official CWD test result indicates the RPLN was independently examined using ELISA and/or IHC testing at Colorado State University (see Schwabenlander et al. 2021, citation 26 in main text). ND=Not Detected

| Animal ID | Age | Sex | Official CWD Test Results for RPLN | MN-QuIC Results |
|---|---|---|---|---|
| 197190 | Female | Yearling | Positive | Positive |
| 197270 | Female | Fawn | ND | ND |
| 197273 | Female | Adult | ND | ND |
| 197264 | Female | Adult | ND | ND |
| 197266 | Female | Adult | ND | ND |
| 197265 | Female | Adult | ND | ND |
| 197267 | Male | Fawn | ND | ND |
| 197268 | Female | Fawn | ND | ND |
| 197269 | Female | Adult | ND | ND |
| 197271 | Female | Adult | ND | ND |
| 197272 | Male | Fawn | ND | ND |
| 197225 | Male | Yearling | Positive | Positive |
| 197065 | Male | Adult | Positive | Positive |

**Supplementary Table 3. Metadata of Examined Wild White-Tailed Deer Tonsils.** MNPRO = Minnesota Center for Prion Research and Outreach. RPLN=medial retropharyngeal lymph node. Official CWD test result indicates the RPLN was independently examined using ELISA and/or IHC testing at Colorado State University (see Schwabenlander et al. 2021, citation 26 in main text). ND=Not Detected

| Process ID | Animal MNPRO ID | Age | Sex | Official CWD Test Results for RPLN | RT-QuIC Results for Tonsil | MN-QuIC Results |
|---|---|---|---|---|---|---|
| 000734 | 000166 | Adult | Female | Positive | Positive | Positive |
| 000735 | 000197 | Yearling | Male | Positive | Positive | Positive |
| 000736 | 000250 | Adult | Male | Positive | Positive | Positive |
| 000737 | 000333 | Adult | Female | Positive | Positive | Positive |
| 000738 | 000360 | Adult | Female | Positive | Positive | Positive |
| 000739 | 000363 | Adult | Female | Positive | Positive | Positive |
| 000740 | 000376 | Adult | Female | Positive | Positive | Positive |
| 000741 | 000384 | Adult | Female | Positive | Positive | Positive |
| 000742 | 000498 | Yearling | Male | ND | ND | ND |
| 000743 | 000508 | Adult | Male | Positive | Positive | Positive |
| 000744 | 000735 | Adult | Female | Positive | Positive | Positive |
| 000745 | 000065 | Adult | Female | ND | ND | ND |
| 000746 | 000067 | Adult | Female | ND | ND | ND |
| 000747 | 000091 | Adult | Female | ND | ND | ND |
| 000748 | 000093 | Adult | Female | ND | ND | ND |
| 000749 | 000230 | Fawn | Female | ND | ND | ND |
| 000750 | 000232 | Adult | Female | ND | ND | ND |
| 000751 | 000283 | Fawn | Male | ND | ND | ND |
| 000752 | 000415 | Fawn | Female | ND | ND | ND |
| 000753 | 000431 | Adult | Male | ND | ND | ND |

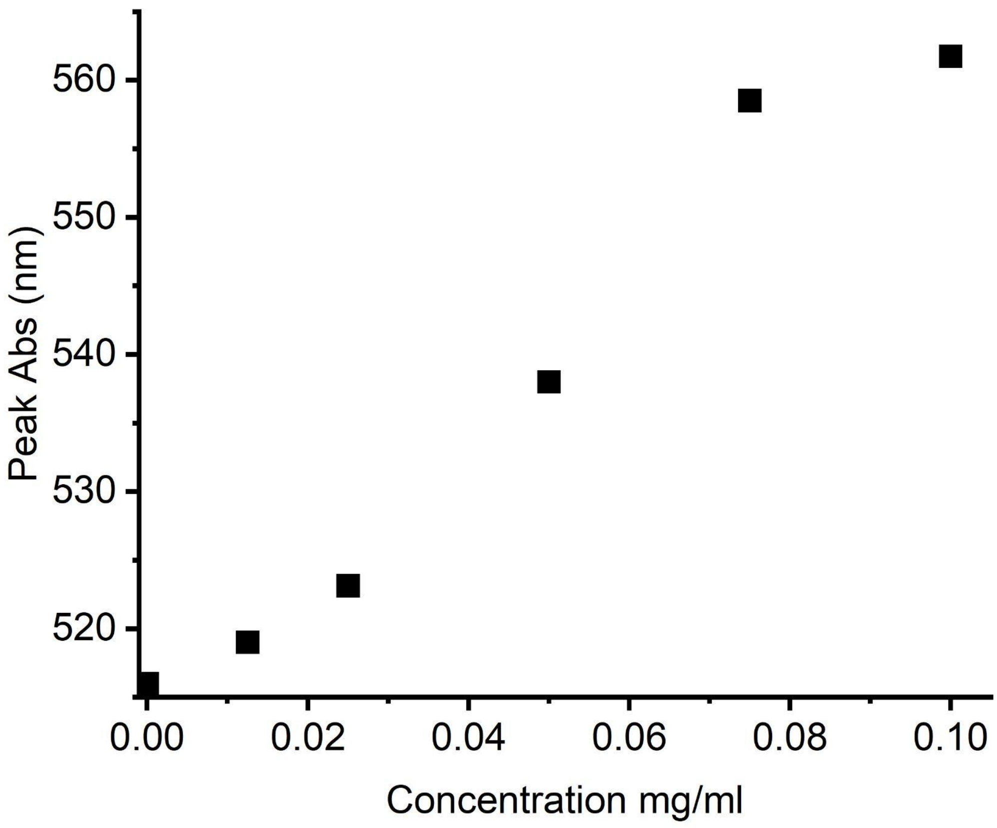


**Supplementary Figure 2.** Wavelength of peak absorbance of AuNPs spiked with various concentrations of native HaPrP substrate in standard mastermix buffer.

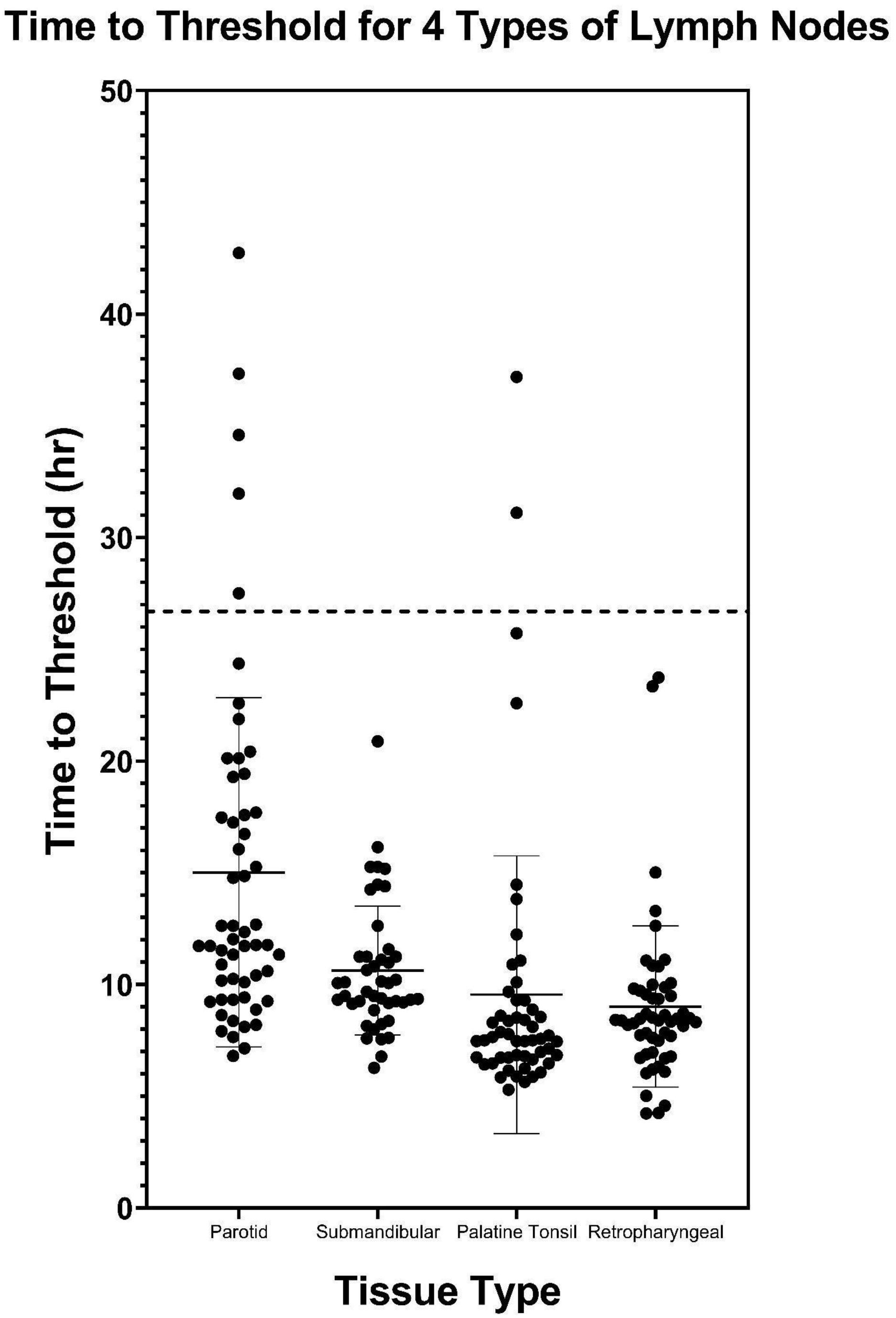


**Supplementary Figure 3:** Prion seeding activity as shown by time (in hours) to Thioflavin-T fluorescence threshold via RT-QuIC reactions for various CWD positive white-tailed deer lymph tissues. Dashed line identifies a 24hr threshold used for MN-QuIC analyses of retropharyngeal lymph nodes.
